# Homeostatic plasticity rules that compensate for cell size are susceptible to channel deletion

**DOI:** 10.1101/753608

**Authors:** Srinivas Gorur-Shandilya, Eve Marder, Timothy O’Leary

**Author notes:** **For correspondence:** (TOL).

## Abstract

Neurons can increase in size dramatically during growth. In many species neurons must preserve their intrinsic dynamics and physiological function across several length scales. For example, neurons in crustacean central pattern generators generate similar activity patterns despite multiple-fold increases in their size and changes in morphology. This scale invariance hints at regulation mechanisms that compensate for size changes by somehow altering membrane currents. Using conductance-based neuron models, we asked whether simple activity-dependent feedback can maintain intrinsic voltage dynamics in a neuron as its size is varied. Despite relying only on a single sensor that measures time-averaged intracellular calcium as a proxy for activity, we found that this regulation mechanism could regulate conductance densities of ion channels, and was robust to changes in the size of the neuron. By mapping changes in cell size onto perturbations in the space of conductance densities of all channels, we show how robustness to size change coexists with sensitivity to perturbations that alter the ratios of maximum conductances of different ion channel types. Our findings suggest that biological regulation that is optimized for coping with expected perturbations such as size changes will be vulnerable to other kinds of perturbations such as channel deletions.

## Introduction

All animals grow, and increase in size over time. In some animals, the physical size of neurons and neural circuits grows along with the body size of the animal, yet continuously maintain their function. For example, action potential frequency and amplitude in Retzius neurons in the leech is constant over four-fold increase in cell diameter (*De-La-Rosa Tovar et al., 2016*). The A4I1 neuron in locust shows similar wind-sensitivity in first instar nymphs and in adults, despite a greater than three-fold increase in cell size (*Bucher and P2üger, 2000*). RPeD1, a neuron in the respiratory central pattern generator in *Lymnaea*, maintains resting membrane potential, spike amplitude and membrane resistance despite two-fold growth from juvenile to adult (*McComb et al., 2003*). In crickets, patterns of motor neuron output responsible for song production appear up to four moults before the adult form (*Bentley and Hoy, 1970*). In lobsters, the co-ordinated bursting activity of pyloric neurons is indistinguishable between juveniles and adults, despite a many-fold increase in their size (*Bucher et al., 2005*). Even in embryonic stages, when these cells are even smaller, the pyloric circuit central pattern generator (CPG) expresses spontaneous albeit less stereotyped rhythmic activity (*Casasnovas and Meyrand, 1995*; *Richards et al., 1999*).

Neurons achieve their target behavior by expressing specific combinations of voltage and calcium gated ion channels (*Prinz et al., 2003*). This leads to the question of how neurons maintain a target function by regulating ion channel expression as the size of the cell increases. Can regulatory mechanisms preserve neuron behavior during growth?

Homeostatic regulation has been proposed as a mechanism that can tune neuronal parameters like ion channel densities to drive neuronal activity to some desired set point (*LeMasson et al., 1993*; *Davis, 2006*; *Liu et al., 1998*; *Turrigiano and Nelson, 2004*; *Turrigiano et al., 1994*; *Turrigiano, 2007*). Changes in neuron size that alter the activity of the neuron may therefore allow activity-dependent homeostatic mechanisms to compensate for the perturbing size change (*Davis and Bezprozvanny, 2001*). Activity dependent feedback control of membrane current expression can therefore potentially compensate for size changes without requiring a mechanism to explicitly measure cell size.

We investigated this hypothesis using simplified biophysical models of pacemaker neurons that use a single sensor of calcium concentration to control ion channel expression homeostatically. Previous work showed that this class of self-tuning models recapitulates experimental observations such a correlations in ion channel density (*Schulz et al., 2006, 2007*; *Temporal et al., 2014*; *Garcia et al., 2018*; *O’Leary et al., 2013, 2014*). Furthermore, these models provide robustness to some forms of perturbations in membrane conductances, but can be sensitive to other perturbations like channel deletions (*O’Leary et al., 2014*). This property is important, because in spite of widespread evidence for homeostatic compensation in the nervous system, there are clearly situations when such mechanisms can fail and potentially be the source of a pathology (*O’Leary, 2018*). We asked under what conditions such simple homeostatic regulation can compensate for changes in cell size, and why some perturbations like channel deletions can lead to pathological rather than remedial compensation.

To answer these questions, we constructed single-compartment conductance-based models of neurons that generated specific behaviour that corresponds to neuron types found in invertebrate CPGs. These single-compartment models allowed us to model a dominant feature of neuron growth, the increase in the cell’s surface area and effective volume, but neglect other features like the extension and branching of neurites and other processes.

We show that a single physiological readout of activity, the time-averaged calcium concentration, can compensate for size changes that would otherwise disrupt electrophysiological behavior. By analyzing how size changes perturb electrical properties, we construct a way to map perturbations that simple, activity-dependent channel regulation can compensate for as well as showing how and why certain perturbations cause this kind of regulation to fail. We suggest that biological regulation that is optimized for coping with expected perturbations such as size changes will be vulnerable to other kinds of perturbations such as channel deletions.

## Methods and Materials

### Neuron model

We used a simple single-compartment conductance-based neuronal model with a single membrane potential *V*, that evolves according to

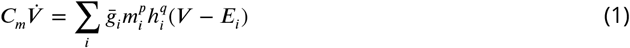

where *C*_*m*_ is the specific membrane capacitance of the cell, 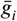 is the maximal conductance density, *E*_*i*_ the reversal potential, and *m* and *h* are activation and inactivation variables of ion channel population *i*. We note that these dynamics do not depend on geometrical properties like the area *A* of the cell or the thin-shell volume *η*.

Another important state variable of the neuron is the intracellular calcium concentration [*Ca*^2+^]_*in*_. In neurons, calcium enters the cell through ligand-gated-and voltage-gated calcium-permeable channels, increasing the cytosolic concentration (*Regehr and Tank, 1994*). Extensive intracellular stores can buffer this calcium influx, and can also act as both a source and a sink of calcium (*Simpson et al., 1995*). Following earlier work that modeled calcium dynamics in neurons (*Buchholtz et al., 1992*; *De Schutter and Smolen, 1998*), we assumed that intracellular calcium concentration evolves according to:

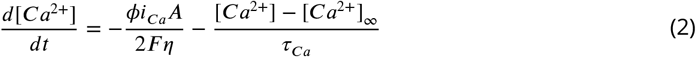

We used a single-compartment conductance-based model of a neuron. The membrane potential of the neuron evolved according to

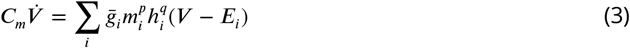

where *C*_*m*_ is the specific membrane capacitance of the cell, 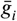 is the maximal conductance density, *E*_*i*_ the reversal potential, and *m* and *h* are activation and inactivation variables of ion channel population *i*. In Figs. 2-4, we used an eight-conductance model with these channels: *A, CaS, CaT, H, Kd, Leak, NaV* and *KCa*. Gating functions for all channels were identical to (*Prinz et al., 2003*). Following earlier work that modeled calcium dynamics in neurons (*Buchholtz et al., 1992*; *De Schutter and Smolen, 1998*), we assumed that intracellular calcium concentration evolves according to:

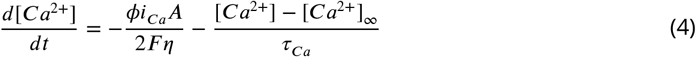

where *ϕ* is a dimensionless parameter, *i*_*Ca*_ is the total calcium current density, *A* is the surface area of the cell, *η* is the volume of the cell available for calcium influx, *τ*_*Ca*_ is the buffering timescale for calcium, [*Ca*^2+^]_∞_ is the resting intracellular calcium concentration and *F* is the Faraday constant. The first term represents calcium influx through voltage-gated calcium channels and the second term represents buffering by intracellular stores, diffusion, and extrusion out of the neuron. The size of the neuron directly affects this equation as both the area of the cell *A* and the volume relevant to calcium influx *η* appear in the first term. Furthermore, the lumped timescale of calcium buffering *τ*_*Ca*_ also depends on the size of the neuron.

### Regulation model

We used a simple model of homeostatic regulation in which transcription and translation depend on the deviation of time-averaged intracellular calcium from some target (*O’Leary et al., 2014*). The concentration of mRNA that encodes channel protein *i* is given by

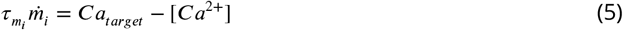

where 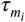 is the timescale of transcription and *Ca*_*target*_ is the average intracellular calcium concentration during target dynamics. Transcription and degradation affect the conductance levels of each channel type according to

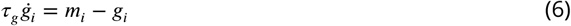

For a given model with some set of conductance densities 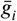, we picked regulation parameters so that

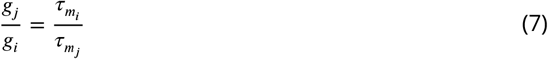

for all pairs *i, j*. Following earlier work (*O’Leary et al., 2014*), we set *τ*_*g*_ to be 5 seconds for all models. While this timescale is probably much longer in reality, we found that this value, which is larger than the slowest timescale of the voltage dynamics, was sufficiently large, and sped up simulations. With these parameters, this homeostatic control scheme could maintain baseline dynamics.

### Growth model

In Figure 1, we dynamically change the size of the neuron using either a linear or exponential growth model. In the linear growth model, the area and the thin-shell volume of the cell change at a constant linear rate 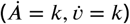, while in the exponential model, 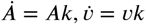.

**Figure 1.**
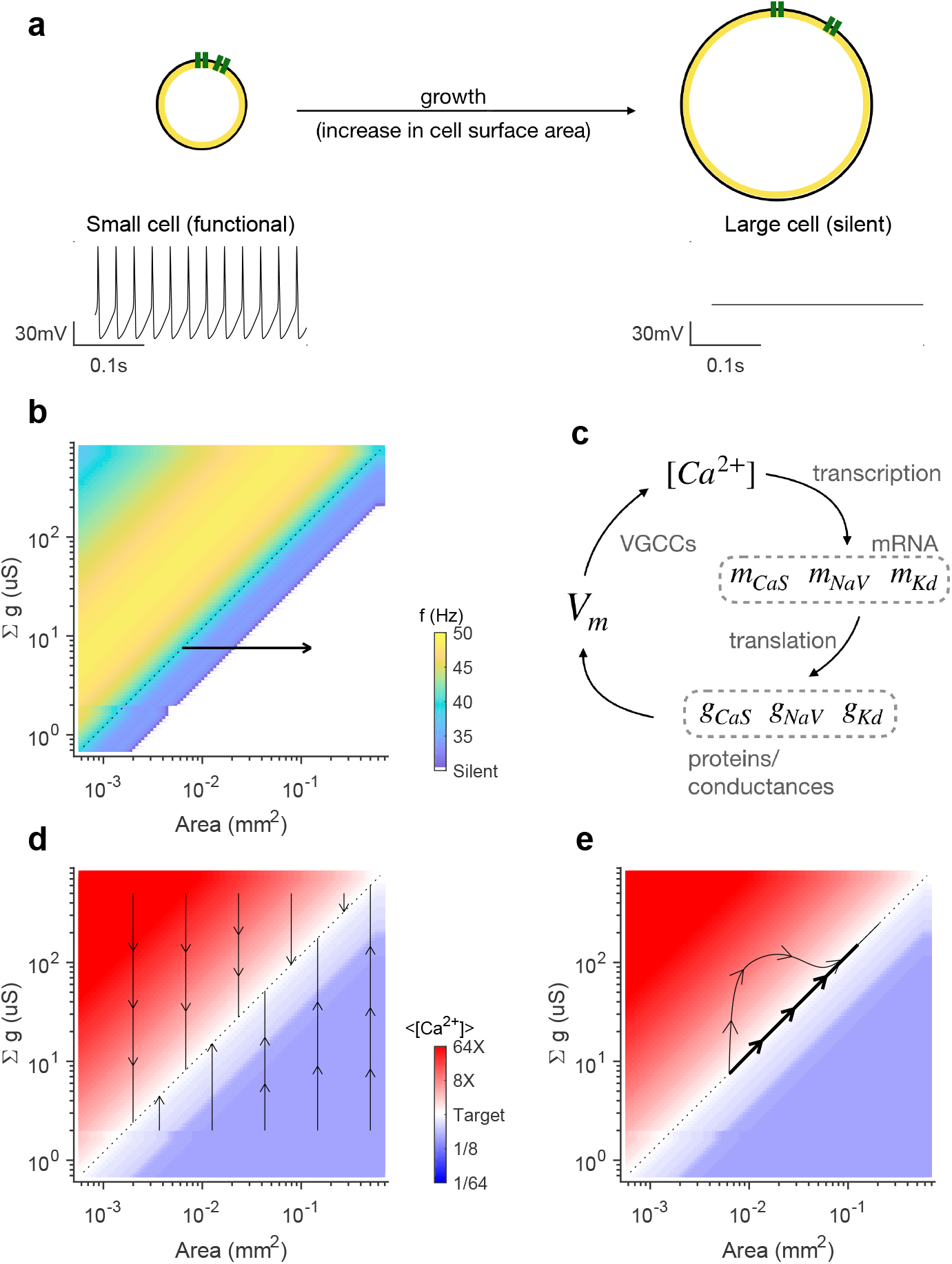
Changes in cell size can disrupt neuron dynamics, but can be compensated for by a mechanism that does not explicitly measure cell size. (a) Growth can cause a tonically spiking cell to become quiescent. (b) Firing rate (color code) of the neuron in (a) as a function of cell size and channel conductance. Arrow indicates growth simulated in (a). (c) A simple biologically-plausible scheme to regulate channel conductances. Transcription of genes coding for channel proteins depends on intracellular calcium concentration, which in turn determines protein abundances, which affect membrane potential, which feed back onto intracellular calcium concentrations through voltage-gated calcium channels. (d) This mechanism makes the line of constant conductance density (dotted line) globally attractive. Arrows show trajectories towards the diagonal under the influence of homeostatic regulation. (e) Regulation of channel conductances during constant linear (thin line) and exponential (thick line) growth. Color codes in (d) and (e) are the same. In all plots, the diagonal dotted line corresponds to constant conductance density.

### Analytical calculation of silent state

A neuron that is silent despite homeostatic regulation must satisfy the constraint that its average intracellular calcium levels are equal to the homeostatic target, *<* [*Ca*^2+^] *>*= *Ca*_*target*_. While in principle this condition can be met while having small oscillations in both *V* and [*Ca*^2+^], we make the simplifying assumption here that the silent state corresponds to fixed points in the calcium and voltage dynamics:

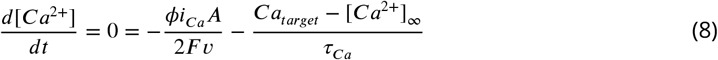

This allows us to solve for *i*_*Ca*_. While there is no closed form expression for *V* at which this equation is satisfied, we can easily calculate it using

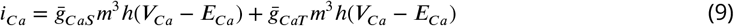

since *CaS* and *CaT* are the only conductances that flux [*Ca*^2+^] into the cell. Here, *m* and *h* can be replaced by their steady state values *m*_∞_ and *h* _∞_ that are simply functions of *V*, since we assume that 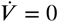. Here *V*_*Ca*_ is the value of the membrane potential where the calcium ODE has a zero.

Similarly, since we know 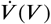(using Eq. 1), we can solve for *V*_*V*_, the value of the membrane potential where the voltage ODE has a zero. In addition, we can compute the marginal stability of this fixed point

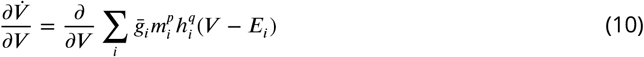

Now we impose the condition *V*_*V*_ = *V*_*Ca*_ at every point in the space of conductance densities and find points where this equation is satisfied.

### Computing calcium level sets and basins of attraction

We defined a ‘calcium level set’ as the set of all points along the plane where the time-averaged intracellular calcium concentration was equal to that of the original model. This level set is concep-tually significant since it is independent of the kinetic parameters of the homeostatic regulation mechanism; yet is the set of points along which the dynamics of the homeostatic mechanism is quenched (since the average intracellular calcium concentration is equal to the target).

In principle, this level set can be found using brute-force methods where a model is simulated at every point along the plane (as in Fig. 2 – Figure Supplement 1). A more eZcient way to find this level set is to adaptively sample the plane, increasing the density of sample points at regions close to the level set. In practice, we did this by initially randomly sampling a few points in the plane, and building a Delaunay triangulation of sampled points. A new point was chosen for sampling based on the largest triangle that contained at least one node above the target calcium level and at least one node below. This process was iterated till an acceptable mesh size was achieved. A similar sampling algorithm was used to find the basins of attraction of the four categories of voltage dynamics when homeostasis was allowed to change the conductance densities of the neuron after perturbation.

### Parameterizing generic high-dimensional perturbations

To characterize generic perturbations in conductance density that are not constrained to a plane (as in Figure 4), we measured the mean and standard deviation of all conductance densities as follows:

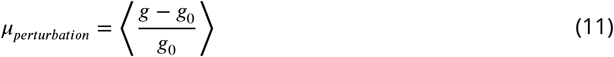

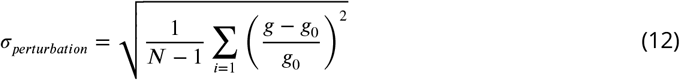

### Building a database of neuron models with similar dynamics

Following earlier work that generated databases of neuron models (*Prinz et al., 2003*), we constructed a set of neuron models with similar intrinsic dynamics by randomly sampling points from a hypercube in the 8-dimensional space of conductance densities. For each point, a model was initialized using these maximal conductances and its intrinsic voltage dynamics checked to see if it fell within acceptable tolerances of a pre-defined target activity. We have made a toolbox to eZciently construct these databases in parallel publicly available at https://github.com/sg-s/neuron-db.

### Model implementation and code availability

All models were implemented in xolotl, a freely available neuron and network simulator (*Gorur-Shandilya et al., 2018*). Voltage and calcium equations were numerically integrated using the exponential Euler method (*Dayan and Abbott, 2001*) using a time step of 0.05 ms.

Code to reproduce all figures in this paper is available at https://github.com/marderlab/size-compensation.

## Results

### Single-sensor homeostatic regulation can compensate for changes in cell size

The model we use in this paper (see Methods) uses a single compartment with one variable to model the membrane potential *V*, and one to model the intracellular calcium concentration. We make the simplifying assumption that the effective volume *η* scales with the area of the cell *A*. This approximation, referred to as the “thin-shell assumption”, is based on the following findings. First, it is known that calcium buffering and diffusion create “micro-domains” of elevated intracellular calcium around calcium channels, suggesting that calcium influx does not lead to large increases in the calcium concentration across the entire volume of the neuron (*Regehr and Tank, 1994*; *Parekh, 2008*; *Tadross et al., 2013*). Second, an important method by which intracellular calcium affects neuron dynamics is by activating calcium-activated potassium (KCa) channels, which are known to localize close to calcium channels (*Fakler and Adelman, 2008*; *Guéguinou et al., 2014*; *Chad and Eckert, 1984*). Third, fitting this model to experimental data revealed that the fast time courses of calcium-sensitive currents could only be reproduced by using a much smaller volume than the geometrical volume, consistent with the idea that the volume accessible to varying calcium is a thin shell within the membrane (*Buchholtz et al., 1992*).

Neglecting the effect of cell size on *τ*_*Ca*_ for the time being, we define “growth” to mean an increase in the surface area *A* and thin-shell volume *η*. Because we assume that *A* and *η* scale together, Eq. 2 is invariant with cell size, and the sole effect of cell growth is a co-ordinated decrease in the maximal conductance density 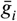 for all channel types in Eq. 1. We found that such an increase in the size of a cell typically leads to a silencing of intrinsic voltage dynamics, since the right-hand side of Eq. 1 is quenched. Figure 1a shows a spiking neuron that stops spiking as its cell size is increased.

Eq. 1 suggests that one solution to maintaining voltage dynamics is to keep the set of conductance densities of all channels constant. This would mean scaling the maximal conductance of all ion channel types with the area of the cell. Fig. 1b shows the firing rate of this neuron as a function of area of the cell and the total maximal conductance of all channels. Left to right motion in this space, corresponding to an increase in cell size shown in Fig. 1(a), leads to a decrease in firing rate and eventually leads to silence. However, firing rate is constant along diagonal lines, which correspond to lines of constant conductance density.

Can a homeostatic feedback mechanism steer a neuron along one line of constant conductance density? Fig. 1c shows a schematic of a simple biophysically plausible regulation mechanism. We assumed that the primary determinants of the neuron’s intrinsic activity were the number of ion channels of each type (sodium channels, potassium channels, and so on) and that the only effect of homeostatic feedback was to regulate the maximal conductance of all ion channel types. We hypothesized that intracellular calcium concentration controls the slow timescale turnover of ion channels. Ion channels control membrane potential dynamics, which in turn feeds back onto intracellular calcium concentration through voltage-gated calcium channels. Following earlier work (*O’Leary et al., 2014*), we used a simple set of equations (see Methods) to model this feedback loop.

where 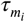 is the timescale of transcription and *Ca*_*target*_ is the average intracellular calcium concentration during target dynamics, and transcription and degradation of ion channel proteins are modeled as relaxing exponentially to the mRNA levels of that channel. This homeostatic mechanism drives the neuron towards realizing a target intracellular calcium concentration, *Ca*_*target*′_, which is set to the average intracellular calcium concentration in the reference model.

We then calculated the average calcium concentration in all points in this space (Fig.1d). Calcium levels are equal to the target level along the line of constant conductance density, suggesting that in this case calcium levels are a good proxy for neuron activity. This regulatory scheme makes the line of constant conductance density globally attractive (Fig.1d), indicating that this scheme can tune conductance densities *without* explicitly needing to measure cell area.

To test these predictions we modeled linear (Fig. 1e, thin line) and exponential Fig. 1e thick line) growth using a fixed rate. Homeostatic regulation was able to steer the neuron onto the line of target intracellular calcium, and thus the line of constant conductance density, and thus the line of desired firing rate, for both linear and exponential growth. Note that in the linear growth model, the change in cell size initially overwhelms the compensatory mechanism, which leads to a transient loss in target dynamics. However, in the exponential growth model, motion is entirely along the diagonal, and neuron dynamics is preserved at all points during growth. Intuitively, this implies that cell growth and channel expression rates need to be approximately matched biologically to maintain physiological function.

### Robustness to perturbations analogous to size change results in sensitivity to some channel-specific perturbations

We have shown that homeostatic regulation can compensate for changes in cell size, but it remains unclear why it can sometimes fail when channels are deleted or over-expressed (*Liu et al., 1998*; *O’Leary et al., 2014*; *Kulik et al., 2019*; *Temporal et al., 2014*). Neurons’ acute sensitivity to perturbations can vary; and homeostatic regulation can compensate for a perturbation, or compensation can be pathological, where perturbations that the neuron is innately robust to can trigger homeostasis to alter voltage dynamics in an undesirable manner (*O’Leary et al., 2014*).

In our framework, size changes are equivalent to a particular form of perturbation in the maximal conductances of ion channel populations, where all conductances are scaled together. A generic perturbation in a neuron with *n* ion channel types is a *n*-dimensional vector, and perturbations like channel deletions are typically incongruent with size changes. This suggests that homeostatic regulation may be good at compensating for perturbations in some directions, but not others.

To quantitatively test this prediction, we analyzed response to perturbations, including size changes, in an 8-conductance neuron model that can exhibit a rich variety of voltage dynamics (*Liu et al., 1998*; *Prinz et al., 2003*), including tonic spiking and bursting.

Understanding the dependence of this homeostatic feedback mechanism in an 8-dimensional space is challenging. Due to the dependence of the feedback mechanism on intracellular calcium, we reasoned that the 8 ion channel types in this model could be grouped into two sets based on whether they directly affected intracellular calcium or not (Fig. 2a). Using a simple classification scheme to group neuron behaviors into one of four non-overlapping classes (Fig. 2b), we then projected the 8-dimensional space of conductance densities onto a two-dimensional plane (Fig. 2c). Regular bursting dynamics with burst periods and duty cycle within ten percent of the target neuron were classified as “canonical” (green), bursting dynamics outside this range were classified as “other bursting” (purple). The other two states were a silent state (blue) and a tonically spiking state (red).

**Figure 2.**
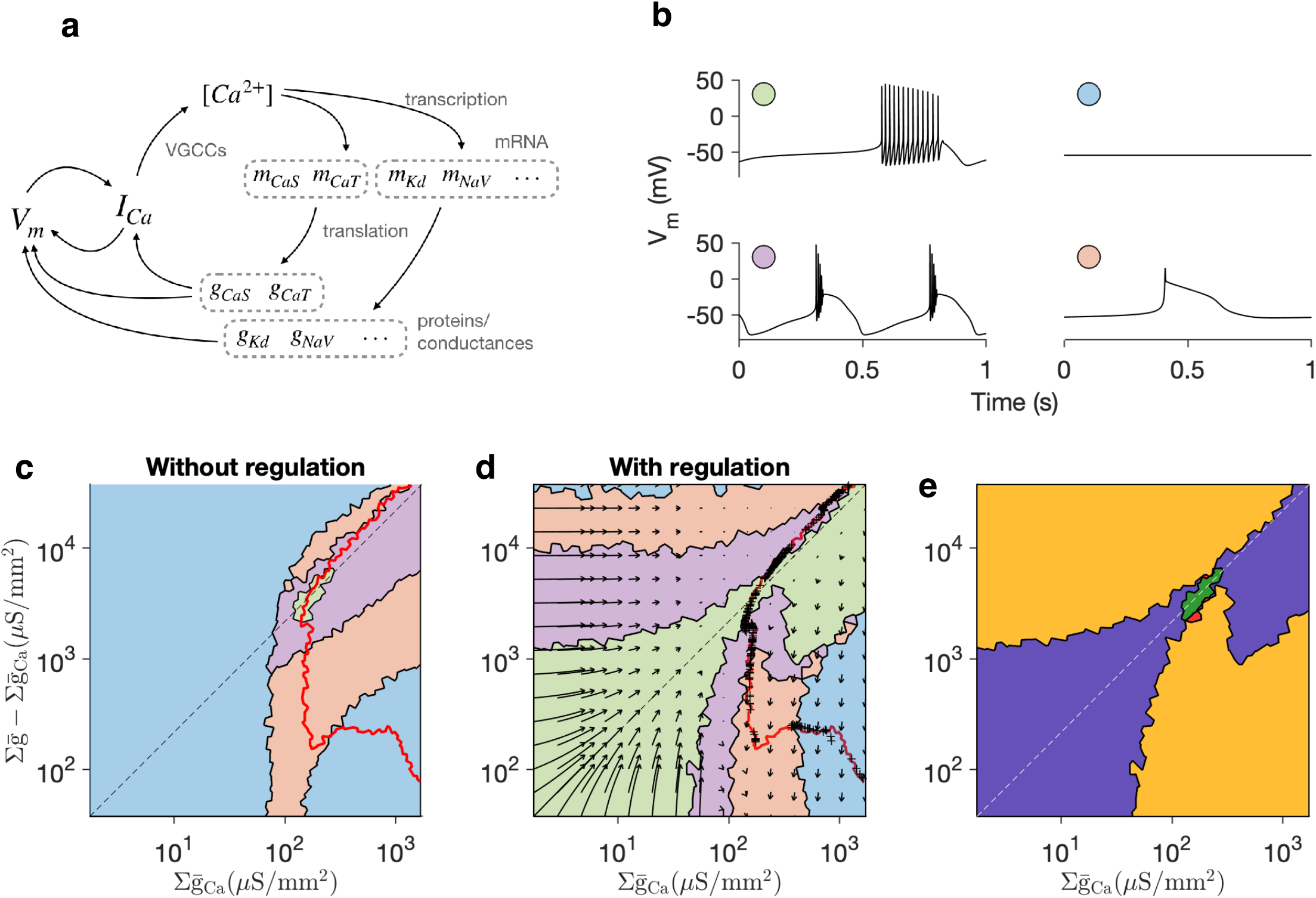
Ratio-preserving regulation rules confer robustness to size changes, at the cost of sensitivity to orthogonal perturbations. (a) Feedback loop segregated into variables that directly affect intracellular calcium and those that do not. (b) Four classes of neuron dynamics as conductance densities are varied: canonical (green), silent (blue), non-canonical bursting (purple) and one-spike bursting (pink). (c) Distribution of these dynamical classes across space of conductance densities. The diagonal corresponds to changes in cell size. The red line joins points in this space where the average intracellular calcium is equal to the average of the reference model. Along the red line, the regulation mechanism, no matter what its kinetic parameters, will be inactive. (d) Basins of attraction of different dynamical states with homeostatic regulation. Cells perturbed along the diagonal (corresponding to a change in size) return to the reference point in conductance space. Perturbations that are further from the diagonal lead to other states. (e) Segmentation of space based on response to perturbation and compensation: robust to perturbation, compensation restorative (green); sensitive to perturbation, compensation restorative (blue); sensitive to perturbation, compensation pathological (orange); robust to perturbation, compensation pathological (red).

We then calculated a level set of calcium where average intracellular calcium levels were equal to that of the target model (Fig. 2c, red line). This line passed through all four states, suggesting that many different states could result from homeostatic compensation, even for neuron models with different parameters that displayed spiking dynamics (Fig. 2– Figure Supplement 1).

To determine which perturbations could be recovered from, we numerically calculated the basins of attraction of these four dynamical states by initializing models at different points in the plane (Fig. 2d). For example, all perturbations that move the conductance densities of the neuron within the green basin can be compensated from, because homeostatic regulation drives the conductances back to the target state, and the neuron eventually generates its original voltage dynamics. There was a total overlap between the basin of attraction of the target state and the diagonal, suggesting that all size change perturbations can be compensated for perfectly. Strikingly, the basin of attraction of the target state was relatively broader close to the origin and relatively narrower at the reference state. This suggests that a larger relative de-correlation in ion channel conductance densities can be compensated for at low absolute values of ion channel densities, as would be seen in a developing neuron that had begun to express its ion channels.

Finally, the calcium level set was an excellent predictor of asymptotic conductance densities of neurons perturbed to start from all over the plane (black crosses on red line, (Fig. 2c)). This is surprising since the motion of the system is not restricted to this two-dimensional plane, but can instead exist in the full 8-dimensional space of conductance densities. The correspondence between the calcium level set (calculated without knowledge of the kinetic parameters of the homeostatic regulation) and the asymptotic states, suggests that trajectories tend to preserve ratios of conductance densities.

Robustness to perturbations along the diagonal (corresponding to changes in cell size) coexisted with sensitivity to off-diagonal perturbations. Mapping the flow field in this plane (Fig. 2d) reveals that flows close to the diagonal are restorative and drive the neuron to the original set of conductance densities, while flows far from the diagonal can drive the neuron to homeostatic targets far from the reference model. Thus, perturbations in a direction orthogonal to the diagonal can lead to a very different steady-state outcome, resulting in pathological compensation.

We constructed a map of the sensitivity of feedback regulation to perturbations (Fig. 2e). Only a small region corresponded to acute robustness to perturbation (Fig. 2e green). A much larger region, along the diagonal, corresponded to sensitivity to perturbation but where homeostasis could compensate for the perturbation (Fig. 2e blue). Intriguingly, regions where the neuron was acutely robust to perturbations existed close to the reference model, where compensation was pathological, and drove the neuron to states with undesirable voltage dynamics (Fig. 2e red). These regions, though close to the reference model, did not lie on the diagonal. This suggests that ratio-preserving regulation rules, which may have evolved biologically to confer robustness to size changes, are vulnerable to perturbations in the expression of some, but not all channel types.

### Predicting failure of homeostatic compensation

In the previous sections, we showed that that the calcium level sets correspond not only to the desired physiological bursting dynamics, but also other bursting dynamics, tonic spiking, and silence. The existence of these regions of parameter space are necessary, but not sufficient, for a pathological homeostatic outcome because the error of the feedback signal in these regions is zero.

To find sufficient conditions for failure of a calcium-dependent feedback mechanism, we focussed on the region in parameter space where calcium level sets coincided with silence. While previous work had also suggested that homeostatic compensation to perturbations such as channel deletions could render neurons silent, it remains unclear, mechanistically, why some perturbations can be compensated for, and why some render the neuron silent, and cannot be recovered from. Periods of silence have been observed during “crashes” when experimental perturbations exceed the permissive range, and are typically beyond neurons’ ability to compensate for (*Haley et al., 2018*; *Haddad and Marder, 2018*; *Tang et al., 2012*).

We hypothesized that homeostatic rules can trap neurons in silent states when the membrane potential is sufficiently depolarized so that a constant influx of calcium through calcium channels occurs due to window currents. We assumed that the silent state corresponds to quenched dynamics in both voltage and calcium. Because the calcium level is constrained to be the same as that of the reference model (the calcium target), we can solve for the fixed points of Eq. 2 to obtain the voltages at which the calcium dynamics are quenched. We note that voltages at which these fixed points occur do not depend on channels that are not calcium channels, since they do not affect calcium dynamics. Fixed points in the calcium ODE exist only for large values of the calcium channels conductance density (Fig. 3a), suggesting that silent states that cannot be recovered from are impossible below a critical calcium channel conductance density.

**Figure 3.**
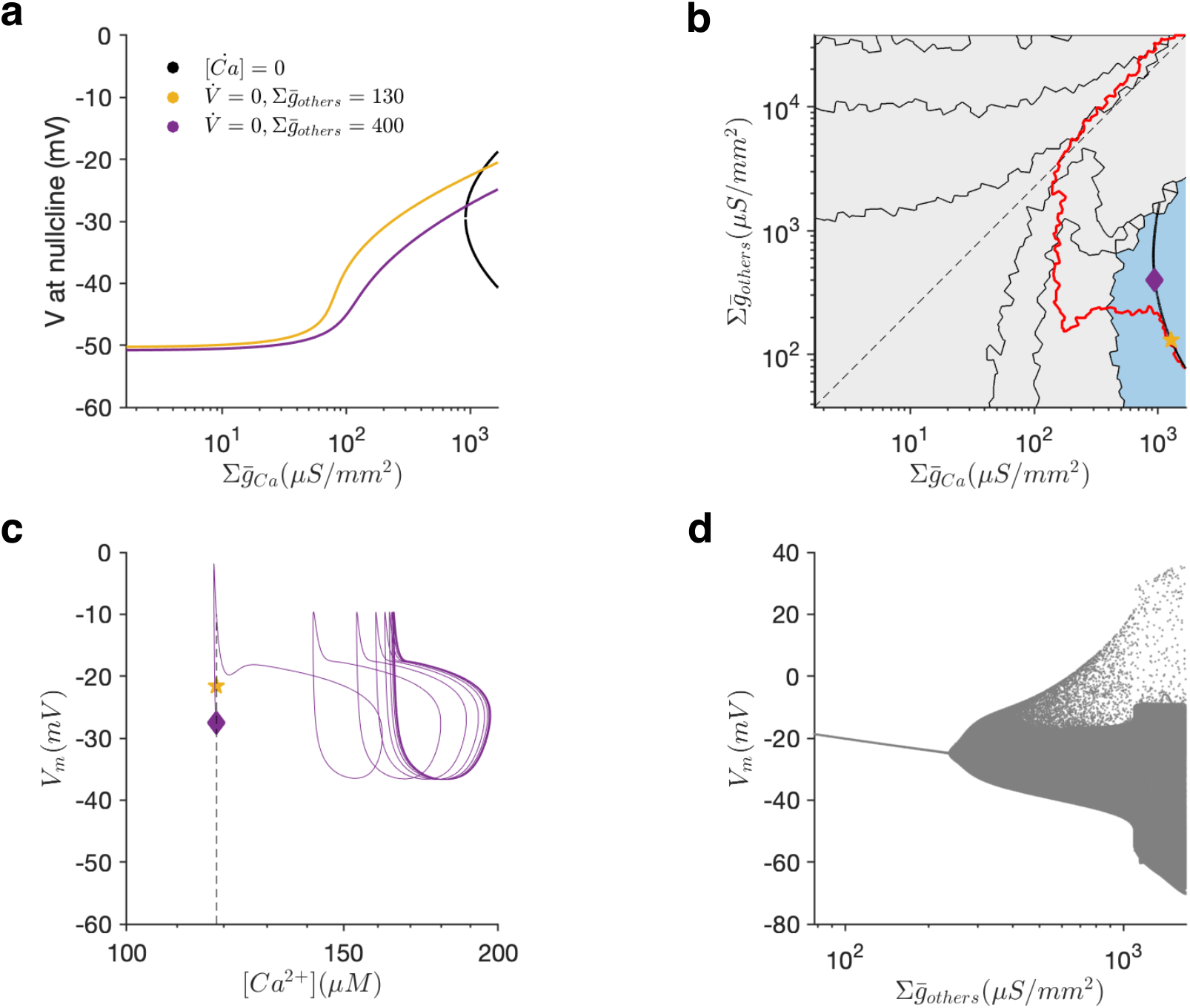
Analytical prediction of when perturbations can silence the neuron. (a) Values of membrane potential corresponding to zeros in the calcium or voltage ODE. The intersection of colored and black curves corresponds to fixed points in the neuron dynamics. (b) Segmentation of conductance density space highlighting the basin of attraction of the silent state (blue). Red line is the calcium level set computed by numerically integrating the model. Black line is the analytical prediction of the location of silent solutions. Pentagram (yellow) and diamond (purple) indicate two locations along the analytically predicted silent state that we shall examine more closely. (c) Dynamics of a neuron model without homeostasis when initialized at these two points. The state indicated by the pentagram (yellow) is stable, but the state indicated by the diamond (purple) is not. Dashed line indicates target calcium of homeostatic regulation. (d) Membrane potential of the neuron along the analytical solution for the silent state, parameterized by the Y-axis in (b). At a critical point, the quiescent state is destabilized, corresponding to the point where the analytical solution and the calcium level set diverge in (b).

Similarly, we can solve for fixed points in the voltage ODE (Eq. 1). The intersection of these two sets of curves corresponds to fixed points in both the voltage and calcium dynamics (Fig. 3a). Plotting the location of points in parameter space where both fixed points overlap reveals a smooth line (black) that partially overlaps with the calcium level set (red), and is entirely contained in the numerically computed basin of stability of the silent state (Fig. 3b).

Why does the analytically calculated set of silent states contain a branch that does not overlap with the numerically computed calcium level set? One possibility is that while all points in the analytical set are indeed silent, their stability properties change along the curve. The voltage fixed points are always unstable, and the upper branch of the calcium fixed points are always stable, suggesting that the overall stability of the silent state may depend on which dynamics dominates as a function of position along the set. To test this, we examined two points along the analytical set, one where it coincided with the numerically measured calcium level set (Fig. 3b, yellow star), and one on the branch that diverged from the numerically measured calcium level set (Fig. 3b, purple diamond). While a neuron initialized at the yellow star remained quiescent (Fig. 3c), a neuron initialized at the purple diamond spontaneously left the silent state, and settled on a periodic sub-threshold orbit (Fig. 3c). Plotting the voltage as the parameters of the neuron model are varied along the analytically calculated set reveal that the neuron switches from a silence to sub-threshold oscillations to spiking (Fig. 3d), which is caused not by the regulatory mechanism but by a destabilization of the fixed point in the intrinsic voltage and calcium dynamics of the neuron.

Together, these results show that simple, calcium-dependent channel regulation mechanisms can be inherently sensitive to channel deletions and produce pathological compensation, even for perturbations that may not affect neural behaviour acutely.

### Robustness to scale perturbations persists across projections and neuron models

Up to this point, we have analyzed the effect of perturbations using a specific projection of the high-dimensional space of conductances. The dimension of the full space of conductance densities is equal to the number of distinct ion channel populations *N*, and the projections shown in the preceding sections do not capture the full space. Similarly, the full level set of calcium is also high dimensional, since it exists in the full *N*-dimensional space, and appears as lines in these projections only because intersections with the projection plane are plotted. The projection chosen in the preceding sections emphasized the distinct contribution of calcium currents, leading to the question if the general features seen hold true for other projections.

We repeated the perturbation analysis for two additional perturbations (Fig. 4a-b). No matter what projection is chosen, the diagonal always corresponds to a change in size, since along that line all conductance densities are scaled together. For both additional projections, we found that the diagonal is entirely contained in the basin of attraction of the canonical state (Fig. 4a-b, green zone) and that the calcium level set intersects with the diagonal exactly once. Taken together, these results suggest that scale perturbations (size changes) can typically be compensated for by homeostasis, but off-diagonal perturbations (e.g. channel deletions, pharmacological manipulations) may not be.

**Figure 4.**
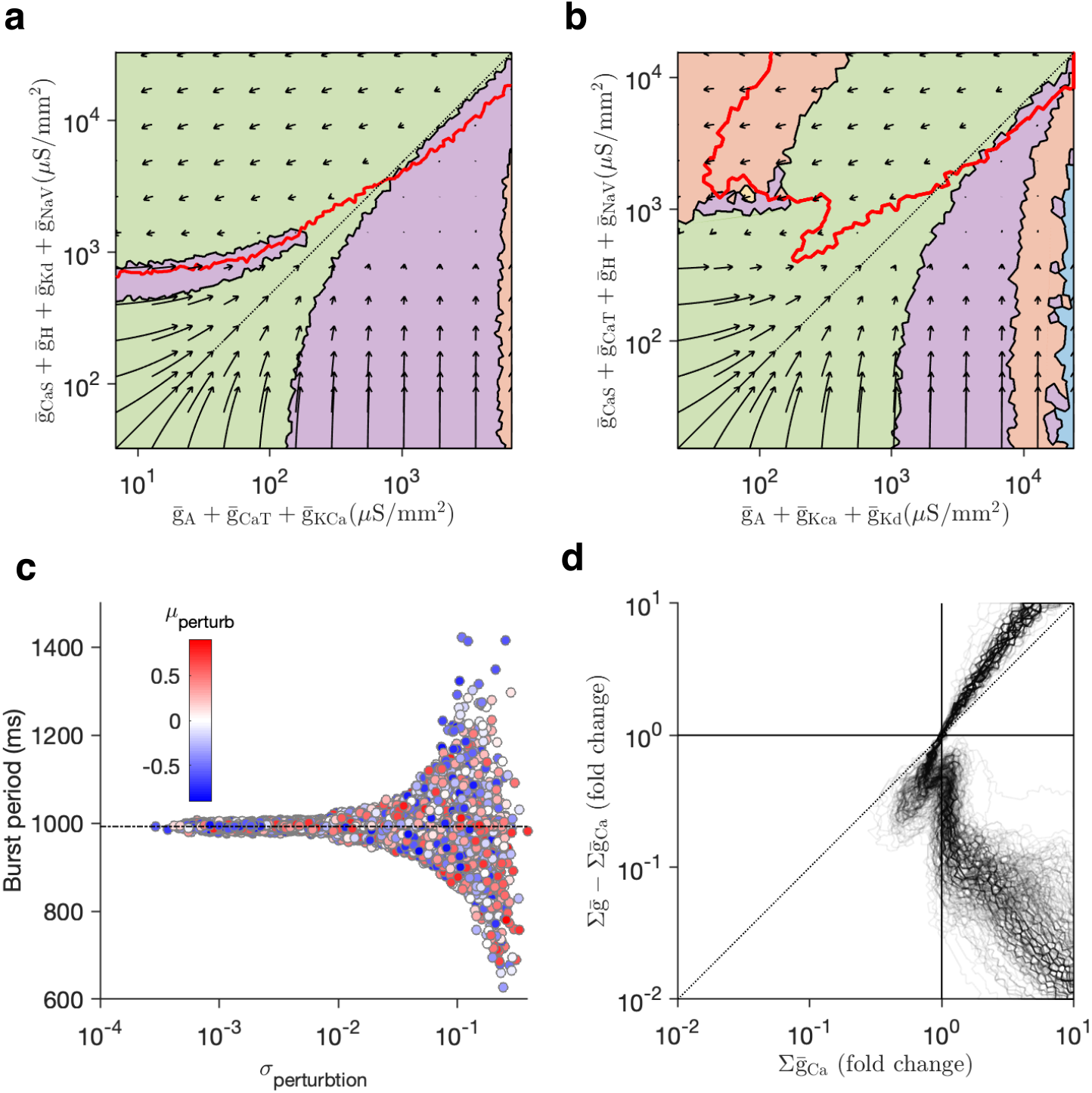
Robustness to scale perturbations persists across projections and neuron models. (a-b) Basins of attractions of target dynamical state (green) and calcium level sets (red lines) for two additional projections. (a) Projection onto arbitrary subsets of conductance densities. (b) Projection onto conductance densities of outward (X) and inward (Y) currents. In both (a) and (b), the diagonal is contained in the basin of attraction of the target state (green), and the diagonal intersects the calcium level set only once. (c) Burst period after homeostatic compensation to perturbations parameterized by mean and standard deviation. (d) calcium level sets for a database of 350 neuron models that share similar voltage dynamics but differ in their conductance parameters. Despite large variation in conductance values and ratios of maximal conductances, in almost all models, the level sets intersect with the diagonal only once.

A generic perturbation in the conductance densities of ion channels in a neuron is high-dimensional, and does not occur on a plane. What effect does a typical perturbation have on the homeostatic system, and what features of the perturbation determine if compensation is restorative or pathological? We parametrized random perturbations in the full-dimensional space by their mean and variance (relative to the conductance densities in the original model, see Methods), and measured the burst period of neuron models after recovery from perturbation (Fig 4c). Consistent with our two-dimensional perturbation analysis, the ability of the homeostatic mechanism to recover from perturbations depended not on the mean value, but on the variance, suggesting that homeostasis could be simultaneously robust to large size changes (low-variance, correlated changes) and sensitive to ratio-disrupting perturbations.

Theoretical and experimental work has shown that many different sets of maximal conductances can lead to similar voltage dynamics (*Prinz et al., 2003*; *Golowasch et al., 2002*; *Goldman et al., 2001*; *Taylor et al., 2009*; *Marder, 2011*; *Caplan et al., 2014*; *Swensen and Bean, 2005*; *Aizenman et al., 2003*). Are all models that share a similar dynamics equally robust to changes in size and off-diagonal perturbations? Picking a regularly bursting neuron with a well-defined burst period and duty cycle, we searched the 8-dimensional space of conductance densities to find 350 sets of maximal conductance densities that displayed similar dynamics, though the variation of individual conductance densities and ratios of conductance densities was significant ((Fig. 4– Figure Supplement 1). For every model, we measured the average intracellular calcium at baseline, and then computed a level set of calcium for every model where the set of calcium was the same as in baseline. Despite the large variation in individual parameters, all calcium level sets were strikingly similar, and almost all level sets intersected with the diagonal exactly once (Fig. 4d). The similarity of calcium level sets across neuron models suggests that different models with diverse maximal conductance densities are robust to changes in size, and can behave similarly to perturbations in their ion channels.

## Discussion

### Consequences of using calcium concentration as a proxy for cell activity

For activity-dependent homeostatic regulation to work, a neuron must be able to measure the deviation of its own activity from some target. A common hypothesis, that we have adopted in this paper, is that neurons can estimate some features of their own dynamics using intracellular calcium concentrations as a proxy for activity (*LeMasson et al., 1993*; *Siegel et al., 1994*; *Gunay and Prinz, 2010*; *Golowasch et al., 1999*; *Davis, 2006*; *O’Leary and Wyllie, 2011*). Because baseline levels of intracellular calcium concentration are very low compared to extracellular levels, depolarization of the membrane that opens calcium channels can lead to a transient calcium influx. Calcium channels therefore can play an important role in directly mapping the voltage dynamics of a neuron onto its intracellular calcium levels. Intriguingly, calcium channels are expressed early in development, suggesting that this part of the feedback loop precedes regulation and expression of ion channels needed for the neuron’s target behavior (*Baccaglini and Spitzer, 1977*; *Liljelund et al., 2000*; *Yamashita and Fukuda, 1993*; *Faure et al., 2001*; *Heusser and Schwappach, 2005*).

However, our results suggest that having too many or too large a fraction of calcium channels can lead to undesirable outcomes, where calcium window currents can fool homeostatic regulation into perpetuating silent states (Fig. 3). Intriguingly, a number of studies of calcium-dependent regulation show that calcium influx through specific channels is required for homeostatic responses, which may make homeostatic regulation more robust biologically (*O’Leary et al., 2010*; *Wheeler et al., 2012*). Developing and growing neurons likely switch regulation rules during their lifetime, using one rule to express channels needed for function and another to maintain ionic conductances within acceptable limits (*Desai et al., 1999*), suggesting that neurons may avoid some forms of pathological compensation by switching regulation rules based on developmental context.

### Consequences of modeling calcium concentration using a single variable

One of the consequences of the simple single-compartment model we have used in this paper is that we use a single variable to describe (a) the calcium concentration in the thin shell that affects the gating of KCa channels, (b) the calcium concentration in the bulk of the soma that likely determines rates of translation, insertion or degradation of ion channel proteins (Eq. 6), and (c) the calcium concentration in the nucleus, that modulates transcription rates (Eq. 5). In real neurons, these three calcium concentrations can likely be very different (*Sala and Hernández-Cruz, 1990*; *De Schutter and Smolen, 1998*), and can have different scaling properties. For example, while the thin-shell calcium concentration can scale with the surface area of the cell, the bulk calcium concentration may scale with the volume of the cell. While this model lacks the details to tease these disparate effects apart, it is useful to consider the biophysical models that are consistent with the assumptions we made to simplify our analysis. The discovery of protein synthesis in neuronal compartments far from the soma, including in dendrites, suggests that local activity-dependent mechanisms can regulate local protein levels (*Steward and Levy, 1982*; *Miller et al., 2002*; *Sutton et al., 2004*). Such mechanisms likely depend on concentration of calcium in some small neighborhood, rather than the calcium concentration in the soma, suggesting that a one-compartment model can mimic some features of local, activity-dependent protein synthesis and degradation (*Ouyang et al., 1999*). Another possibility is that detachment of granules of mRNA at a particular location (*Doyle and Kiebler, 2011*), or the insertion or removal of ion channels into the membrane, can depend on the local calcium concentration. In summary, local activity-dependent regulation mechanisms can be approximated by the model used here, and are an attractive formalism since they do not require co-ordination of sensors and regulators across the neuron.

### Homeostatic compensation of growth in spatially extended neurons is more complex

We examined how neurons could use homeostatic mechanisms to regulate their ion channel conductances to preserve intrinsic dynamics as their size changed in a single compartment model. A single compartment model neglects many of the complexities in real neurons since the neuron is described by a single membrane potential and a single value of intracellular calcium and is assumed to have no internal structural heterogeneity. While single compartment models are clearly only a coarse approximation of real neurons, recent work has suggested that some neurons are surprisingly electronically compact despite their large size and spatially complex morphology (*Otopalik et al., 2017*; *Ray et al., 2019*). Other neurons with long processes that extend their processes as the animal grows (e.g. motor neurons controlling muscles in the extremities) face several additional challenges. First, their dynamics are no longer described by a single membrane potential since they are not electronically compact. Second, channel expression is spatially regulated: for example, sodium channels occur at a higher density on the axon (*Kole et al., 2008*). Third, intracellular calcium levels are not spatially uniform (*Hernández-Cruz et al., 1990*), and can vary substantially along the length of thin processes (*Regehr and Tank, 1994*). Finally, since all mRNA ultimately originates from the cell body, mRNA and ribosomes and other translation machinery, or entire ion channels, must be physically transported from the nucleus to parts of the cell where they are needed (*Doyle and Kiebler, 2011*; *Kosik, 2016*; *Bramham and Wells, 2007*), leading to other regulatory problems with the transport of cargo (*Doyle and Kiebler, 2011*; *Williams et al., 2016*). Nevertheless, our formalism reveals certain fundamental problems that must be addressed even in more detailed analysis that models neurons as spatially extended.

### Stochastic effects can arise from low copy numbers

Following earlier work, we modeled the gating of ion channels as a deterministic process (*Hodgkin and Huxley, 1952*). Even though the gating of an individual ion channel is a stochastic process, this deterministic model approximates well a large number of ion channels, since their individual fluctuations average out (*White et al., 2000*). However, when neurons are sufficiently small, or when conductance densities are sufficiently low, a neuron, or a piece of membrane in a neuron may have only a small number of channels (*Smith, 2002*). In this limit, the stochastic gating of ion channels becomes substantial, and has been shown to qualitatively change the behavior of neurons (*Chow and White, 1996*; *Sengupta et al., 2013*). Another source of stochasticity is in the regulation mechanism since mRNA copy numbers are typically low (*Kosik, 2016*). It is not clear whether homeostatic mechanisms continue to function in these extreme limits, and there is formal theory to show that low copy numbers present regulation problems that cannot be circumvented (*Lestas et al., 2010*). Such effects will introduce perturbations in the physiology of a cell that we have not accounted for explicitly here.

### Model predictions for non-neuronal cells

Our results suggest that cells that generate stereotyped voltage dynamics despite potential changes in size can use a single sensor, for instance intracellular calcium, to regulate abundances of several different protein types that contribute to the voltage dynamics. Neurons are the archetypal electrically active cells, and can realize this negative feedback loop using calcium-dependent transcription, translation, or channel insertion, voltage-gated channels, and calcium channels. Stereotyped electrical activity is widespread in many different cell types, including bacteria (*Masi et al., 2015*; *Kralj et al., 2011*), pancreatic *β*-cells (*Bertram et al., 2010*) and cardiac cells (*Hund and Rudy, 2000*). Bursting oscillations play an integral role in insulin secretion by pancreatic *β*-cells (*Bertram et al., 2010*), and these cells can compensate for genetic deletions of a critical channel K(ATP) channel population by over-expressing other potassium channels (*Yildirim et al., 2017*) in the mouse, but not in humans. Recent work suggests that these cells can use intracellular calcium as a sensor of voltage dynamics in a negative feedback loop to regulate activity (*Yildirim and Bertram, 2017*). Thus, many other cell types may share common features of homeostatic regulation with neurons such as robustness to size changes and sensitivity to some perturbations.

### Other mechanisms of compensation for growth

In this study, we analyzed how neurons can regulate the densities of ion channel populations as they grow. In addition to this, neurons can regulate a number of other properties during growth. The manner in which a neuron grows can have a critical role in shaping neuronal function. In *isometric* growth, the length and diameter of a neuron or neuron component increase by the same factor, which increases input conductance by the square of the growth factor, and changes the passive properties of the neuron. In *iso-electrotonic* growth, the diameters of neuronal processes increase as the square of their increase in length, which increases input conductance by the cube of the growth factor, but leaves passive properties of the neuron unchanged (*Olsen et al., 1996*). Lateral giant neurons in the crayfish grow isometrically, and and become progressively less sensitive to phasic components of inputs, since high-frequency signals are attenuated to a greater extent (*Edwards et al., 1994a*,b). Retzius neurons in the leech can compensate for an increase in size by increasing the membrane resistance of the dendrites (*De-La-Rosa Tovar et al., 2016*).

Compensation of size change of one neuron can also involve other neurons. In pyramidal cells, as dendrites grow, pre-existing synaptic sites become physically or electrically more distant to the soma (*Davis and Bezprozvanny, 2001*; *Stuart and Sakmann, 1995*). Muscle cells in the crayfish neuromuscular junction maintain a constant level of depolarization, despite a 50-fold increase in their size, by a regulated increase in the presynaptic release and quantal size (*Pulver et al., 2004*; *Davis and Bezprozvanny, 2001*; *Lnenicka and Mellon, 1983*). Circuits early in development can express adult-like rhythmic activity, but their activity can be continuously inhibited by descending cells (*Fénelon et al., 1998*; *Le Feuvre et al., 1999*; *Bentley and Hoy, 1970*). Neuromodulators and co-transmitters (*Fénelon et al., 1999*; *Kilman et al., 1999*) can be sequentially released onto neurons during development, which means that the neuromodulatory context a neuron exists in may depend on its size. Homeostatic mechanisms like synaptic scaling that can compensate for changing levels of input can also help a neuron compensate for changes in input resistance that may occur during growth (*Turrigiano, 2012, 2007*).

## Acknowledgments

We gratefully acknowledge helpful conversations with and careful proofreading by Alec Hoyland and Jason Pipkin.

---

**Figure 2-Figure supplement 1.**
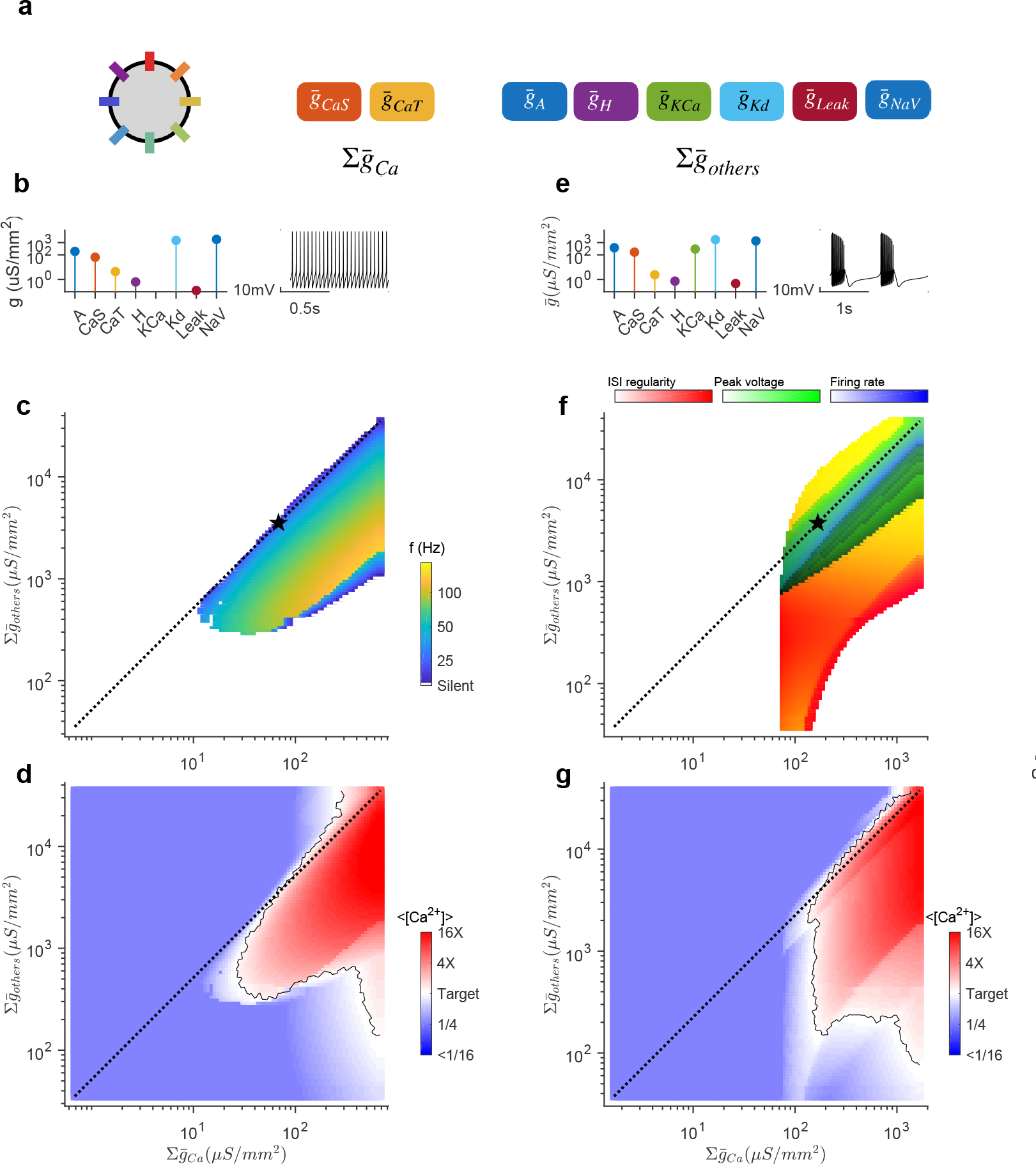
Change in cell size corresponds to a subset of perturbations in the space of conductance densities. (a) A perturbation in the eight-dimensional space of conductance densities in a neuron model can be visualized by projecting onto two dimensions. (b) This combination of conductance densities generates tonic spiking. (c) Firing rate of this model as a function of conductance density. Star indicates the location of the reference model shown in (b). The diagonal preserves all ratios of conductance densities, and therefore corresponds to changes in cell size. (d) Average intracellular calcium as a function of conductance density. Target refers to calcium levels in the reference model. Black line is a level set where calcium levels are equal to the calcium levels in the reference. (e-g) Bursting neuron.

**Figure 4-Figure supplement 1.**
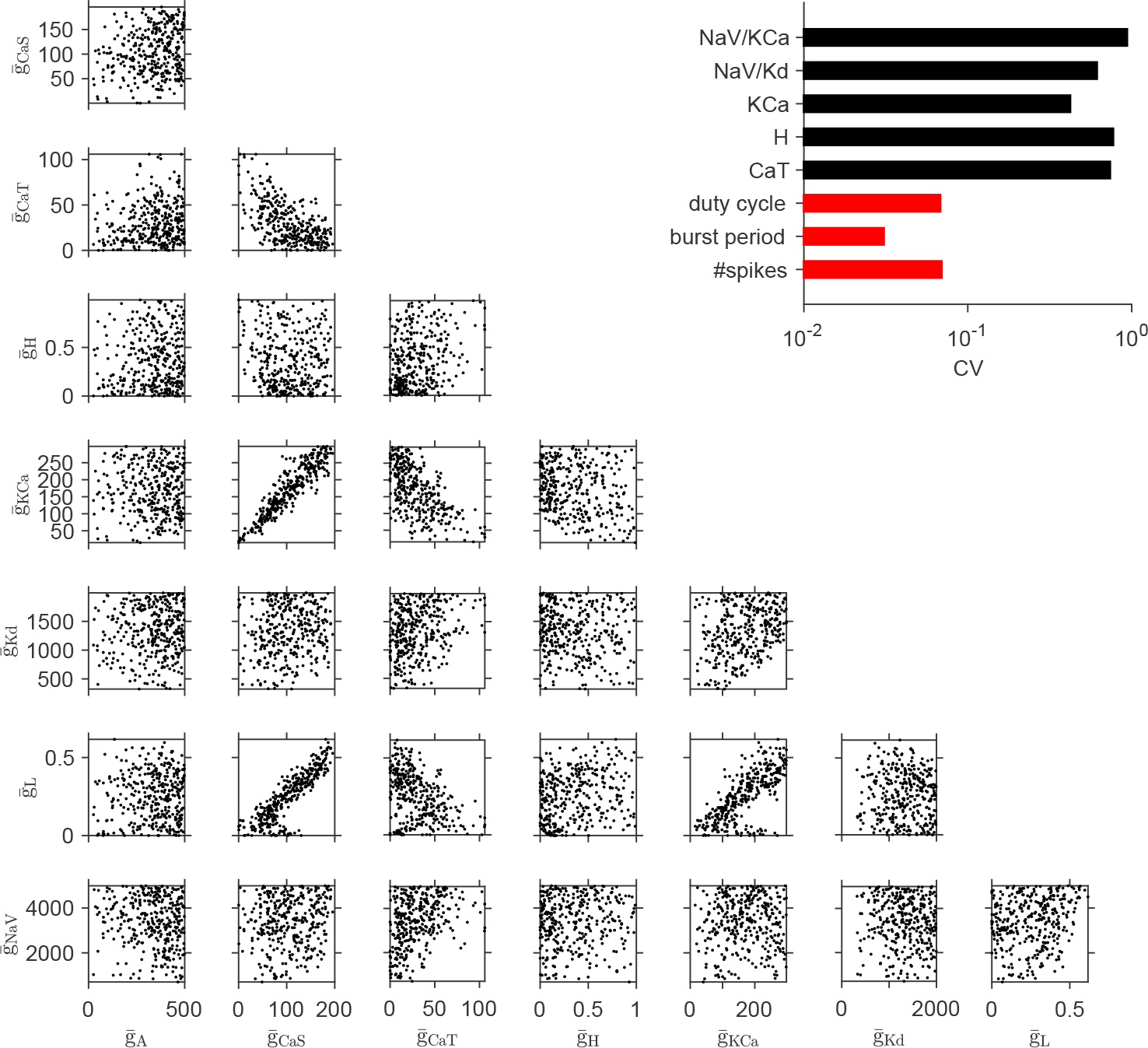
Database of neuron models with varying maximal conductances but similar voltage dynamics. Scatter plots show the pairwise distribution of maximal conductances in a set of randomly sampled neuron models. Each dot in every panel corresponds to a single model. Bar chart compares the coeZcient of variability (CV) between maximal conductances (black) and parameters of neuron voltage dynamics (red). Variability in maximal conductances is much greater than the chosen for tolerance in voltage dynamics. This collection thus constitutes a set of neuron models that display similar voltage dynamics, despite large variation in their parameters.

